# Microtubule-Connexin-43 regulation suppresses arrhythmias and fibrosis in Duchenne muscular dystrophy mice

**DOI:** 10.1101/2022.03.29.486276

**Authors:** Eric Himelman, Julie Nouet, Mauricio A. Lillo, Alexander Chong, Xander H.T. Wehrens, George G. Rodney, Lai-Hua Xie, Natalia Shirokova, Jorge E. Contreras, Diego Fraidenraich

## Abstract

Dilated cardiomyopathy is the leading cause of death in Duchenne muscular dystrophy (DMD) patients due to advancements in skeletal muscle therapies yet limited presence of cardiac treatments. The phosphorylation status of gap junction protein Connexin-43 (Cx43) drives Cx43 remodeling and the development of arrhythmias and fibrosis. Based on evidence that Colchicine drug treatment improves Cx43 phosphorylation and remodeling, we compared the microtubule cytoskeleton in DMD mice (mdx) versus mdx mice genetically altered to be Cx43-phosphorylation-deficient (mdxS3A). Reciprocally, we analyzed the microtubule cytoskeleton in mdx mice genetically altered to be Cx43-phospho-mimicking (mdxS3E). We found a link between the phospho-status of Connexin-43 and regulation of microtubule organization, in which phospho-dead Cx43 (S3A) inhibits improvements seen with Colchicine treatment in mdx mice, and phospho-mimic S3E promotes microtubule reorganization in mdx mice. A reduction in arrhythmias and fibrosis suggests an unsuspecting Cx43-microtubule link for translational corrective activities for DMD cardiomyopathy.

## INTRODUCTION

Duchenne muscular dystrophy (DMD), caused by the loss of function mutation in the DMD gene dystrophin, is the most common and fatal form of muscular dystrophy, affecting 1 in every 5,000 males (1–3). Despite the benefit of therapies directed toward skeletal and respiratory muscle improvement, the lack of effective treatments tailored for DMD cardiac failure has limited DMD patient survival (4). Recent reports detail that dilated cardiomyopathy occurs in up to 95% of DMD individuals 20 years and older (5). To date, there is no cure for DMD and DMD-associated heart failure, limiting treatment strategies to management of symptoms and slowing disease progression (1). Thus, the focus has shifted toward studying principal DMD-specific cardiac pathomechanisms to understand the disease better and develop novel treatments to improve patient outcomes.

Aberrant expression of the cardiac gap junction protein Connexin-43 (Cx43) plays a role in the development of cardiomyopathy in the mdx mouse model of DMD (6–9). Our previous study found that a reduction of phosphorylation of Cx43 C-terminal serine residues S325/S328/S330 triggers pathological Cx43 redistribution to the lateral sides of cardiomyocytes (Cx43 remodeling) in DMD hearts, thereby exacerbating the cardiac phenotype (9). We found that knock-in mdx mice where the Cx43 serine-triplet, replaced with phospho-mimicking glutamic acids (mdxS3E), were resistant to Cx43 remodeling with a corresponding reduction of Cx43 hemichannel activity. Furthermore, mdxS3E mice displayed protection against inducible arrhythmias, the development of cardiomyopathy, and related lethality. On the other hand, knock-in mdx mice with non-phosphorylatable alanines substituted into the Cx43 serine triplet (mdxS3A) exhibited severe cellular and physiological phenotypes much like mdx (9).

Furthermore, we revealed an important link mediating dystrophin loss with subsequent Cx43 remodeling in DMD hearts – the microtubule (MT) cytoskeleton. The dense and disorganized MT network amplifies stretch-induced aberrant Ca^2+^ release triggering ventricular arrhythmias in mdx mice (10, 11). We found that disruption of hyper-densified, disorganized MTs with Colchicine (Colch) treatment protected against Cx43 remodeling in 4-6 month-old mdx mice, concomitant with enhanced S325/S328/S330 phosphorylation (pS-Cx43) (9). However, it is unknown if pS-Cx43 is the key downstream driver of Cx43 localization in dystrophic hearts after microtubule correction. Additionally, even though pS-Cx43 is suggested to promote Cx43 assembly immediately before gap junctional formation (12, 13), its potential role in mediating MT stability and trafficking remains undefined.

Here, we sought to 1) determine the importance of pS-Cx43 in mediating Colch-induced Cx43 remodeling prevention and 2) its role in regulating dystrophic microtubule behavior. We found that Colch treatment in mdxS3A mice, contrary to Colch treatment in mdx mice, does not protect against Cx43 remodeling and associated Cx43 hemichannel activity. Also, the Colch treatment failed to protect against inducible arrhythmias and MT organization in mdxS3A. Finally, we found that mdxS3E mice displayed a normalization in cardiomyocyte MT density, an enhanced Cx43-β-tubulin interaction, and an improvement in microtubule organization and directionality, mimicking WT mice. These changes triggered by S3E led to improvements in cardiac fibrosis and lethality in severe dystrophic mouse models. Together, our findings suggest a reciprocal relationship between pS-Cx43 and MT dynamics in dystrophic hearts that provides a strong foundation for novel therapeutic options in cardiac pathologies, including DMD cardiomyopathy.

## RESULTS

### Colchicine fails to correct Cx43 remodeling in mdxS3A mice

We first sought to assess the impact of Colch-treatment on the dystrophic microtubule (MT) cytoskeleton of mdx mice in which the serine-triplet of Cx43 was replaced with non-phosphorylatable alanines (mdxS3A). Colch (0.4mg/kg) or Saline was administered using implanted mini osmotic pumps for four weeks. We assessed Cx43 localization following Colch treatment by conducting Cx43 immunohistochemistry alongside ID-marker N-Cadherin in ventricular cryosections derived from Colch or Saline treated WT, WTS3A, mdx, and mdxS3A mice. Confocal imaging revealed significant Cx43 remodeling at baseline(9) and after Colch treatment in mdxS3A hearts (Figure 1a). Quantification of relative Cx43 signal found at the IDs confirmed Colch did not improve Cx43 remodeling for mdxS3A, and WTS3A, in opposition with mdx hearts (Figure 1b) (9). To further demonstrate reduction of remodeled Cx43 following Colch, we measured ethidium bromide dye uptake in perfused isolated hearts from mice stressed using Isoproterenol (ISO, 5mg/kg) 30 minutes before isolation to exacerbate hemichannel activity (9, 14–17). Ethidium bromide (ethidium, 5μmol/L) uptake, a hemichannel-specific, plasma membrane-impermeable dye, was significantly reduced in mdx ventricular cryosections while mdxS3A or WTS3A displayed no significant reduction in dye uptake following Colch (Figure S1a-b). These results suggest that Colch treatment fails to prevent remodeling of Cx43 S3A, and S3A may promote remodeling, independent of dystrophin loss.

**Figure 1.**
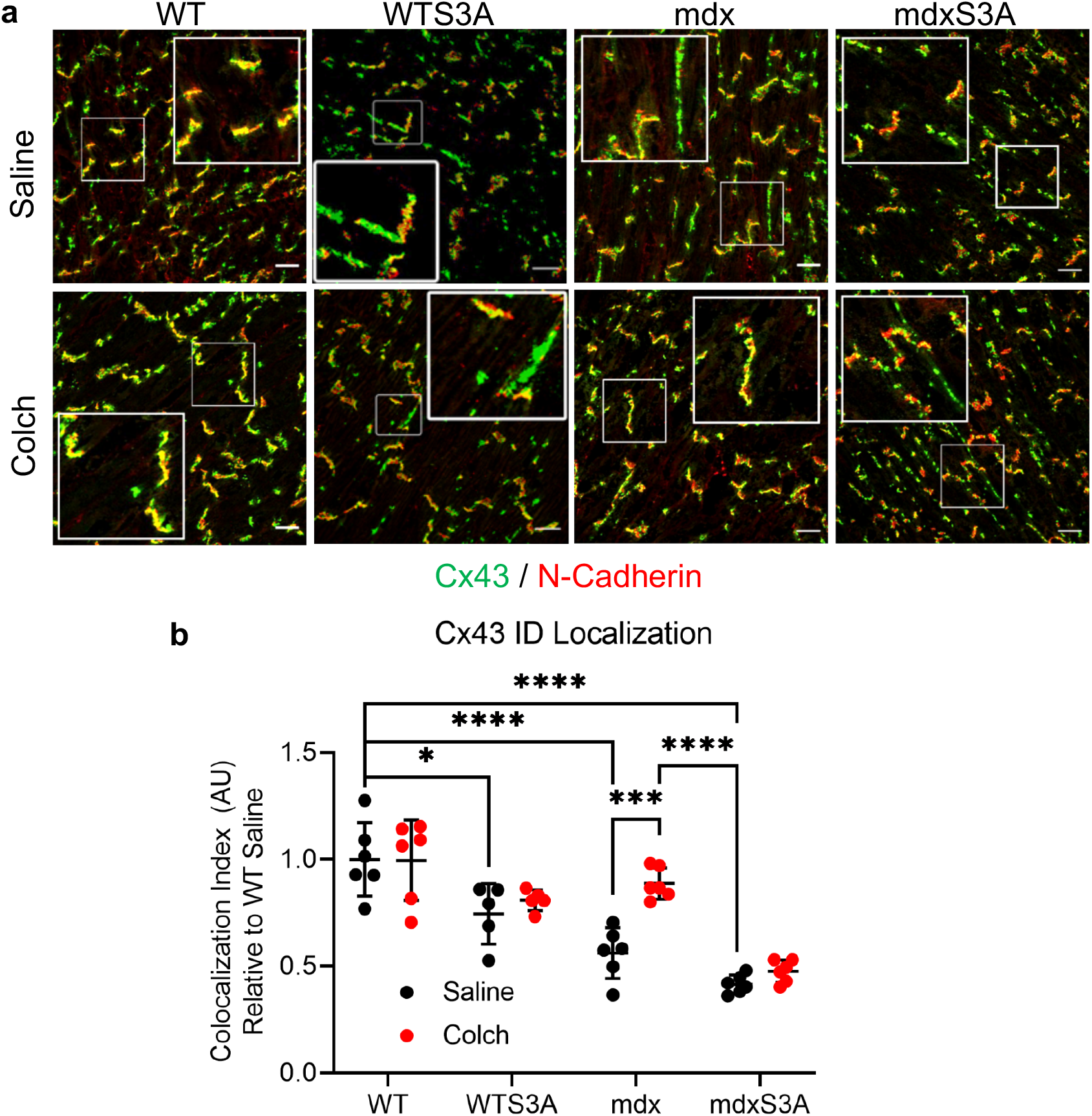
Cx43 remodeling in the hearts of mdxS3A mice following Colch treatment. (**a**) Representative confocal immunofluorescence images, with insets, of cardiac cryosections stained with Cx43(green) and N-Cadherin (red). Scale bar 20μm. (**b**) Quantification of Cx43/N-Cadherin co-localization in confocal immunofluorescent images, processed in Fiji ImageJ. All data points normalized to the WT Saline group mean. *n*=6 (WT ± Colch, mdx ± Colch, mdxS3A ± Colch), *n*= 5 (WTS3A ± Colch). Each dot represents a mean value per mouse: 3-5 images containing 15-20 N-Cadherin positive intercalated discs (IDs) were analyzed per heart. (2-way ANOVA, Tukey’s multiple comparisons test, F(3, 38), ****=p < 0.0001, ***=p < 0.001, *=p < 0.05). Data are presented as means ± SEM.

### Colchicine fails to protect mdxS3A mice from Iso-induced arrhythmias in vivo

To investigate the *in vivo* cardiac conduction phenotype following Colch treatment, we performed a shortterm ECG study in conscious mdx and mdxS3A mice using surgically implanted ECG telemetry devices. Following an initial 60-minute baseline reading, Colch-treated and Saline-treated mice were followed for 72 hours, wherein they received three once-daily ISO (5mg/kg) injections. Following ISO administration, both Saline-treated mdx and mdxS3A and Colch-treated mdxS3A mice exhibited arrhythmogenic events such as pre-ventricular contractions (PVCs) and ventricular tachycardia (VT). In contrast, Colch-treated mdx mice exhibited either single, minor arrhythmic events or none throughout three days of analysis (Figure 2a). When the severity of arrhythmias per day was stratified using an arrhythmia scoring method (9, 18), Colch-treated mdx mice had a significantly reduced arrhythmia score compared to their Saline counterparts, while Colch did not diminish arrhythmia severity in mdxS3A animals (Figure 2b). In addition to arrhythmia severity, we measured a significant reduction of total arrhythmogenic occurrences throughout each 24-hour period in Colch-mdx mice, while no such reduction was observed in Colch-mdxS3A mice (Figure 2c). Together, these results strengthen hypo-phosphorylated Cx43 as a crucial target for arrhythmia protection in DMD mice, particularly following treatments aimed to reduce cardiac stress and normalize the dystrophic microtubule cytoskeleton.

**Figure 2.**
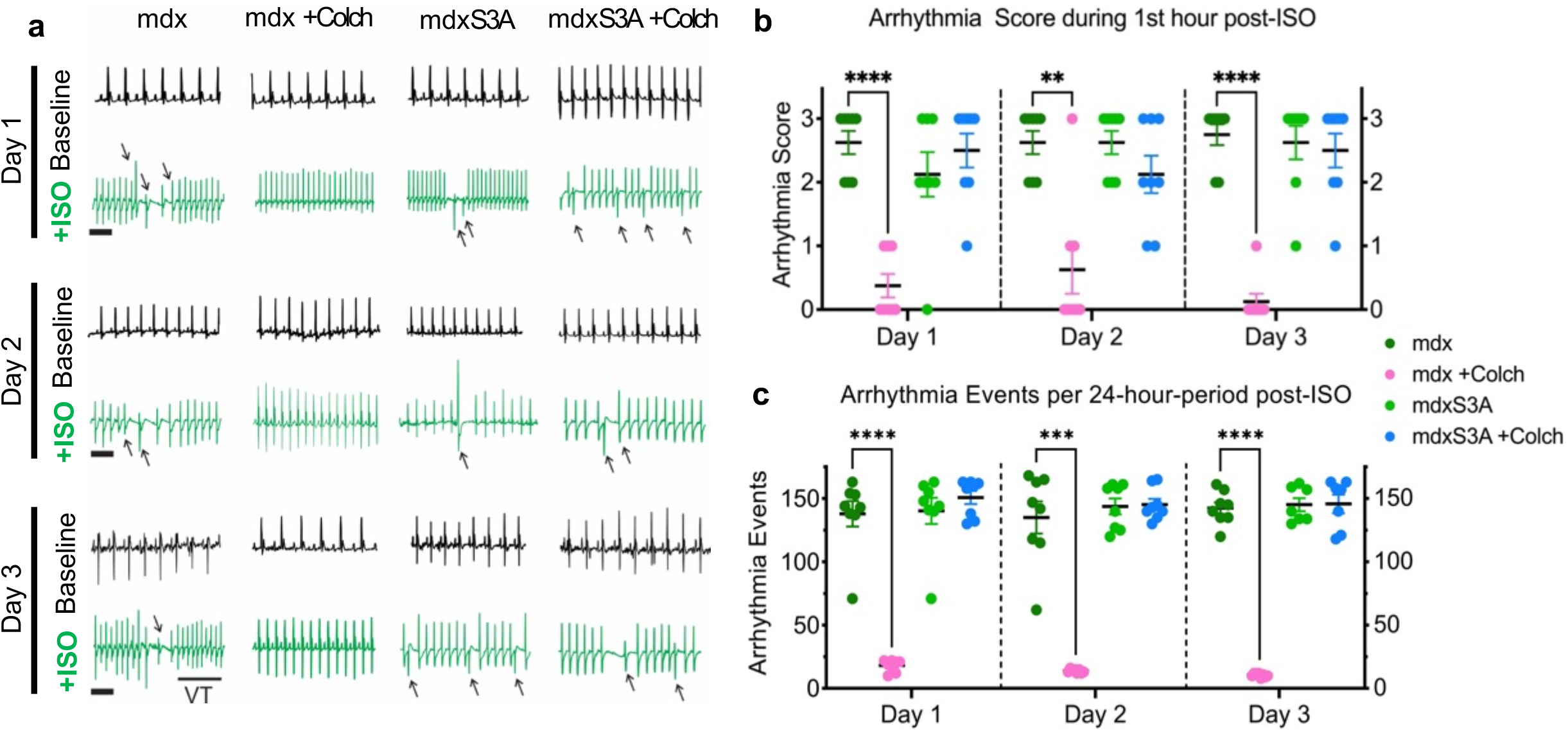
Isoproterenol-induced arrhythmias in mdxS3A mice following Colch treatment. (**a**) Representative ECG recordings obtained from conscious 4-6 month-old mice with implanted telemetry devices. The first two rows represent recordings taken on Day 1 before (Baseline, black) and 1 hour after the first ISO injection (ISO, green). The middle two recordings from Day 2 represent before (Baseline) and 1 hour after the second (24 hours) ISO injection. The last two recordings, Day 3, represent at before (Baseline) and 1 hour after the third (48 hours) ISO injection. Black arrows indicate pre-ventricular contractions (PVCs), consecutive black arrows indicate consecutive PVCs. VT = ventricular tachycardia. Scale bar, 200ms. (**b**) Arrhythmia scores following ISO administration based on a pre-determined scale where 0 = no arrhythmias, 1 = single PVCs, 2 = double PVCs, 3 = non sustained VT (3-10 PVCs), atrioventricular (AV) block on respective days of analysis. (**c**) Quantification of total arrhythmogenic events, including PVCs, double PVCs, VT, or AV block, during the respective day of analysis. (**b-c**) Lab Chart 8 software was used to identify and analyze arrhythmogenic events. *n*=8, all groups. (1-way ANOVA, Tukey’s multiple comparisons test, ****=p < 0.0001, ***=p < 0.001, **=p < 0.01). Data are presented as means ± SEM.

### Colchicine fails to improve MT organization in mdxS3A mice

Because the MT cytoskeleton governs Cx43 trafficking and remodeling(9, 19–22), we sought to study MT dynamics in the dystrophic, Cx43 phospho-dead environment. To evaluate the organization of the MT cytoskeleton, we performed confocal microscopy of Colch treated isolated cardiomyocytes stained with β-tubulin. To analyze the overall organization in the inter-myofibrillar MT network of the cardiomyocyte, we used TeDT analysis (23, 24) (Figure 3c-e). The TeDT analysis demonstrated that mdx and mdxS3A cardiomyocytes displayed a pathological increase in orthogonally oriented MTs, as illustrated by the increased amplitude seen in the magnified region containing orthogonal MTs (60° - 120°, Figure 3D and 88°-92°, Figure 3e)(25). However, Colch-treated mdx myocytes displayed a more organized pattern, much like WT, as evidenced by the confocal images (Figure 3a), density quantifications (Figure 3b), and subsequent TeDT analyses (Figure 3c-e), while these parameters were not rescued in Colch-treated mdxS3A myocytes (Figure 3a-e). The results suggest that Colch fails to organize the microtubule cytoskeleton in myocytes derived from mdxS3A mice.

**Figure 3.**
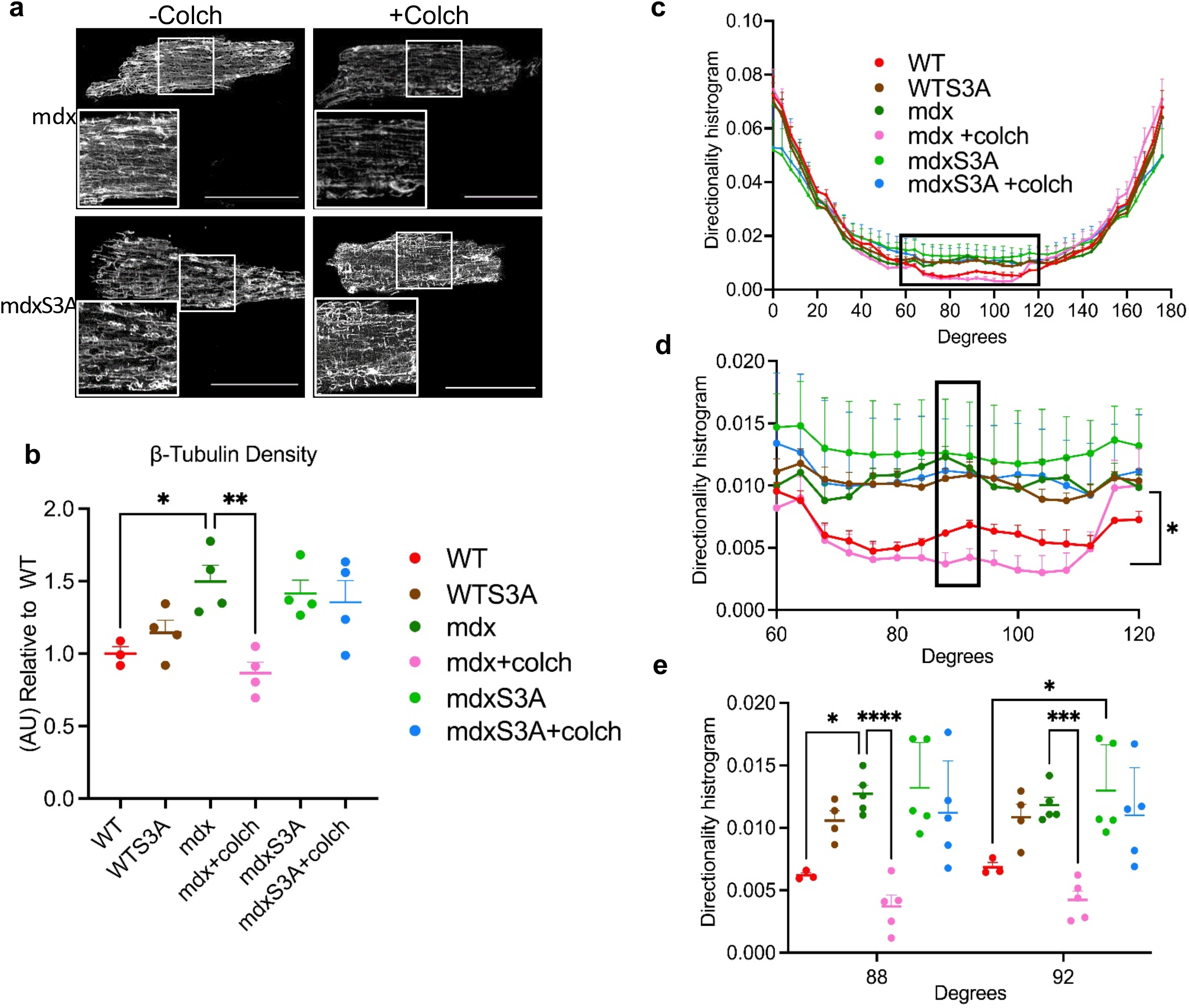
Organized microtubules in cardiomyocytes of mice treated with Colch. (**a**) Representative confocal images of β-tubulin (gray) staining in inter-myofibrillar region of isolated cardiomyocytes derived from 4-6 month-old WT, WTS3A, mdx, mdxS3A with and without Colch treatment. White boxes indicate magnified insets, processed in Adobe Photoshop. Scale bar, 50μm. (**b**) Quantification of β-tubulin immunofluorescent density, processed in Fiji ImageJ. *n*=4, 16 cells per group. (One-way ANOVA, Tukey’s multiple comparisons test, **=p < 0.01, *=p < 0.05.) (**c**) Microtubule directionality determined by TeDT analysis; 0° is defined as the longitudinal axis of the cell, 90° is defined as the orthogonal axis of the cell. (**d**) Magnification of the highlighted region in (**c**). (Two-way ANOVA, F(15, 336), reported as row factor in each comparison *=p < 0.05 mdx versus mdx+Colch.) (**e**) Quantification of directionality histogram values at 88° and 92°, magnified in (**d**). (One-way ANOVA followed, Tukey’s multiple comparisons test (**e**), *** = p < 0.001; *=p < 0.05. *n*=3, n =15 (WT), *n*=4, n=20 (WTS3A); *n*=5, n=20 (mdx±Colch, mdxS3A±Colch) (**b-e**)). Data are presented as means ± SEM.

### Phospho-mimic S3E normalizes dystrophic microtubules

The evidence presented above suggests that interfering with phosphorylation in Cx43 (S3A) promotes pathological remodeling and arrhythmias in DMD mice (Figure 1, S1, 2). The evidence also suggests that the MTs (Figure 3) are pathologically altered in mdxS3A but not in mdx myocytes treated with Colch. This indicates that the correct organization of the MTs is crucial in preventing Cx43 remodeling. However, it is not yet known if mimicking Cx43-S325/S328/S330 phosphorylation (S3E), which itself is anti-arrhythmic in mdx mice (mdxS3E) (9), may also act upon MTs to provide cardioprotection (26, 27). To this end, we studied the organization of the MTs in the hearts of mdxS3E mice. We found that the β-tubulin protein content of mdxS3E hearts was normalized towards WT levels, while mdx and mdxS3A protein levels were both significantly elevated (Figure 4a-b). Supporting our data in Figure 4b, confocal β-tubulin immunofluorescent density was significantly reduced in isolated mdxS3E cardiomyocytes (Figure 4c-d). Mdx and mdxS3A cardiomyocytes displayed a more disorganized MT pattern with a significant increase of MTs oriented orthogonally, as illustrated by the increased amplitude seen when the region containing orthogonal MTs (60° - 120°, Figure 4f and 88°-92°, Figure 4g) was magnified (25). However, mdxS3E myocytes displayed a more organized pattern, much like WT, as evidenced by the confocal images (Figure 4c), density quantifications (Figure 4d), and subsequent TeDT analyses (Figure 4e-g). Together, these data highlight a major role of phospho-Cx43 beyond its previously identified role in remodeling; a role in normalizing the directionality of the MT network. In the DMD heart, where MT derangement is a primary driver of pathological mechano-chemical transduction pathway (24, 26), MT normalization by pS-Cx43 promotion (phospho-mimicking S3E) may provide numerous significant biological improvements in the mechanics of the working DMD cardiomyocyte.

**Figure 4.**
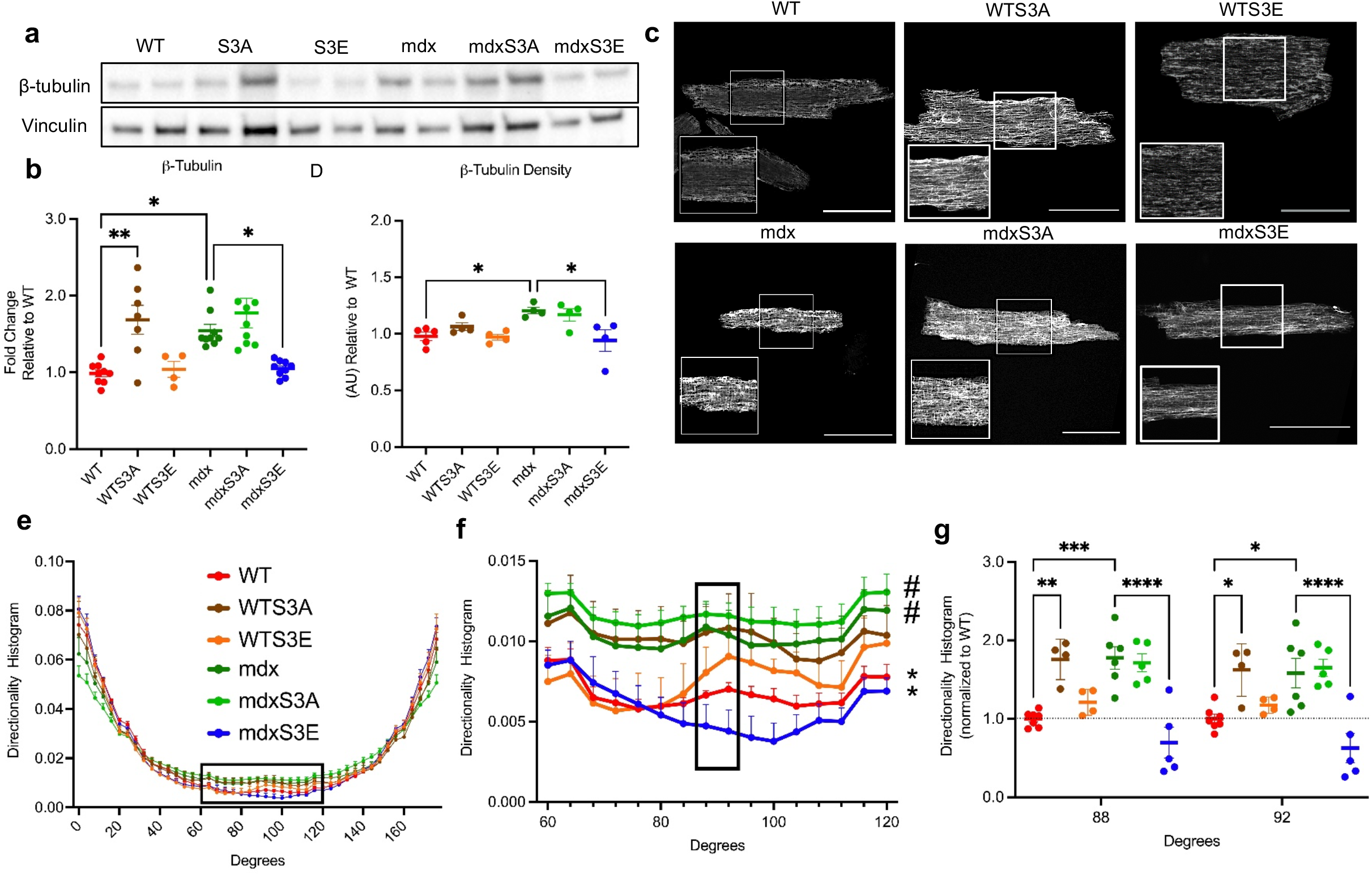
Organized microtubules in cardiomyocytes of mdxS3E mice. (**a**) Representative western blots for β-tubulin, gp91^phox^ and Vinculin (loading control) in lysates from 4-6 month-old WT, mdx, mdxS3A and mdxS3E hearts. (**b**) Quantification of β-tubulin protein levels, normalized to WT protein levels. *n*=4 (WT), *n*=8 (mdx, mdxS3A, mdxS3E). (One-way ANOVA, Tukey’s multiple comparisons test, **=p < 0.01; *=p < 0.05). (**c**) Representative confocal images of β-tubulin (gray) staining in inter-myofibrillar region of isolated cardiomyocytes derived from 4-6 month-old WT, WTS3A, WTS3E, mdx, mdxS3A, and mdxS3E mice. White boxes indicate magnified insets, processed in Adobe Photoshop. Scale bar, 50μm. (**d**) Quantification of β-tubulin immunofluorescent density, processed in Fiji ImageJ. *n*=4, 16 cells per group. (One-way ANOVA, Tukey’s multiple comparisons test, ***=p < 0.001). (**e**) Microtubule directionality determined by TeDT analysis; 0° is defined as the longitudinal axis of the cell, 90° is defined as orthogonal axis of the cell. (**f**) Magnification of the highlighted region in (**e**) (60°-120° of the orientation analysis in (**d**). * = p < 0.05 versus WT, # = p < 0.05 versus mdx. (**g**) Quantification of directionality histogram values at 88° and 92°, magnified in (**f**). (Two-way ANOVA, F(15, 432), reported as row factor in each comparison (**f**) and One-way ANOVA, Tukey’s multiple comparisons test (**g**) ****=p < 0.0001; ***=p < 0.001; **=p < 0.01; *=p < 0.05. N = 6, n = 20 (mdx); N = 5, n = 20 (WT, mdxS3A, mdxS3E) (**e-g**)). Data are presented as means ± SEM.

### Improved Cx43-β-tubulin interaction when pS-Cx43 is improved

Based on the data above and prior evidence indicating Cx43 and β-tubulin as molecular partners (20, 28), we next investigated a possible regulation by direct interaction between Cx43 and β-tubulin in human and mouse dystrophic environments via Cx43 and β-tubulin co-immunoprecipitation assays. In human non-DMD and DMD ventricular lysates, we found increased β-tubulin co-immunoprecipitation in non-DMD tissues relative to levels in DMD tissues (Figure S2a-c). The analysis of mouse tissues shows that the Cx43–β-tubulin interaction was significantly enhanced in mdxS3E compared to mdx ventricular lysates (Figure S2d-f). Similarly, lysates from Colch-treated mdx hearts also displayed a significant increase in β-tubulin co-immunoprecipitation compared to Saline (Figure S 2g-i). These data suggest that a direct mechanism of Cx43-MT trafficking, when improved, may be clinically relevant.

### Phospho-mimic S3E protects against cardiac fibrosis and lethality in severe models of muscular dystrophy

We tested the potency of phospho-mimicking S3E in cardiac pathology development in severe models of DMD mice, Masson trichrome staining was completed to evaluate fibrosis. Utrophin is a dystrophin homologue primarily expressed in the sarcolemma of adult mdx but not WT mice. Utrophin up-regulation in the sarcolemma partially compensates for the absence of dystrophin. The absence of utrophin and dystrophin (mdxutr) leads to a severe phenotype that exacerbates the cardiac pathology followed by premature death between 3-5 months(29–32). The partial absence of utrophin (utr+/-) leads to an intermediate phenotype between mdx and mdxutr^(32, 33)^. We observed that the phospho-mimic mutation S3E led to protection against fibrosis (Figure 5a, b) as well as a protection against premature death (Figure 5c) in mdx mice with partial (mdxutr+/-) or complete absence of utrophin (mdxutr), although in the latter mice the benefits were more modest. Together, these data highlight a modest, protective role for S3E in the development of cardiac fibrosis and against early death in severe models of muscular dystrophy.

**Figure 5.**
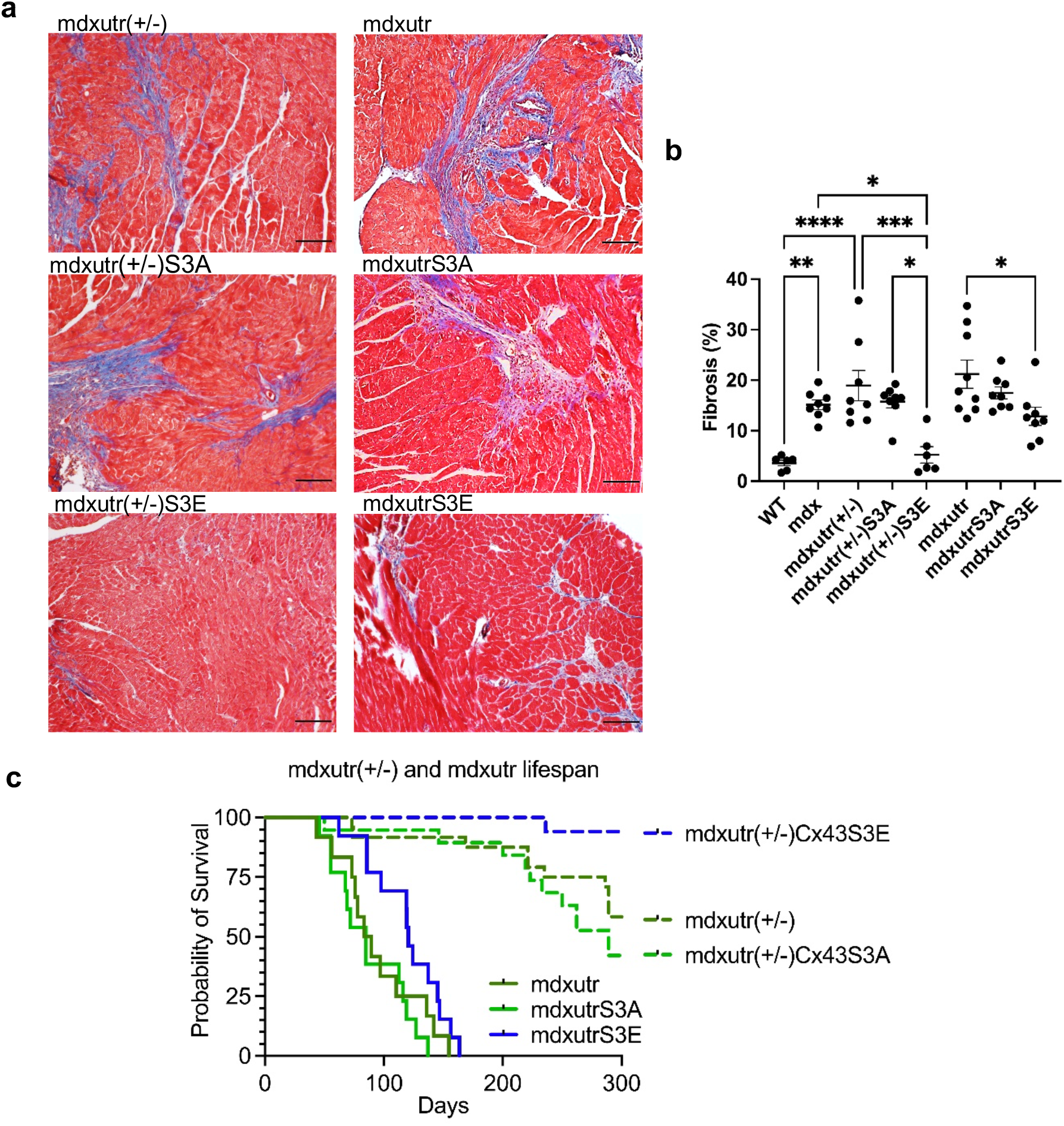
S3E provides protection in more severe models of mdx. (**a**) Representative images of Masson Trichrome histology in hearts of 10-14 month-old mdxutr(+/-), mdxutr(+/-)S3A, mdxutr(+/-)S3E (Left Column) and in 3-4 month-old mdxutr, mdxutrS3A, mdxutrS3E (Right Column). Scale Bar: 200μM. (**b**) Quantification of fibrosis from Masson trichrome histology in mice presented in (**a**). *n*=6 (WT, mdxutr(+/-)S3E) *n*=8 (mdx, mdxutr(+/-), mdxutr(+/-)S3A, mdxutr, mdxutrS3E, mdxutrS3A). (One-way ANOVA, Tukey’s multiple comparisons test, ****=p < 0.0001; ***=p < 0.001; **=p < 0.01; *=p < 0.05) Data are presented as means **±** SEM. (**c**) Kaplan-Meier probability of survival curve for mdxutr, mdxutrS3A, mdxutrS3E, and mdxutr(+/-), mdxutr(+/-)S3A, mdxutr(+/-)S3E. *n* ≥12.

## DISCUSSION

Cardiomyopathy is the leading cause of death in DMD patients (34). The current understanding of DMD cardiomyopathy has increased as of late, but further research is required to reveal important cellular targets to develop more effective cardiac therapies. In prior studies, we identified Cx43 remodeling as an important contributor to DMD cardiac pathology, mainly via hypo-phosphorylation in Cx43 serine residues 325/328/330 (6, 8, 9). We have also shown that the MT cytoskeleton is implicated in the Cx43 remodeling phenotype in DMD hearts (9). These results collectively illustrated the important patho-mechanistic roles of phosphorylation and MT-based trafficking of Cx43 in DMD hearts, but they did not address how these components interface with one another. Furthermore, it is unclear how these mechanisms may intersect with other elements of DMD cardiac disease and the order in which these events may occur *in vivo*, which is ultimately essential when designing new therapies.

While our last report demonstrated negligible differences between mdx and mdxS3A dystrophic cardiac phenotypes (9), this present report illustrates that pS-Cx43 is crucial towards rescuing the cardiac phenotype of mdx mice when the MTs are targeted. Herein, mdxS3A mice, treated with the MT regulator Colch, did not display any protection against Cx43 remodeling, in contrast to what was observed in mdx mice with wild-type Cx43. Accordingly, Colch-treated mdxS3A mice displayed severe cardiac conduction abnormalities following Iso stimulation, while Colch-treated mdx mice were largely protected against Iso-induced arrhythmias. Moreover, Colch-treated mdxS3A failed to normalize organization of the enhanced orthogonal lattice observed in dystrophic MT dynamics. Collectively, the lack of cardioprotection conferred by Colch in mdxS3A mice provides strong evidence indicating that pS-Cx43 is both necessary and sufficient for suppressing Cx43 remodeling. These results strongly suggest that 1) the organization of the MT lattice impacts Cx43 phosphorylation/remodeling, and 2) Cx43 phosphorylation/remodeling impacts the organization of the MT lattice. Thus, rather than MT and pS-Cx43 being organized hierarchically, it appears that MT and pS-Cx43 regulate one another in an amplifying feed-forward loop.

A major finding of our study is that pS-Cx43 itself can normalize MT density and organization in dystrophic hearts, providing important mechanistic information underlying the cardiac protection seen in mdxS3E mice. We argue that in DMD hearts de-phosphorylation and remodeling of Cx43 further disturbs the MTs, and disorganized MTs promotes pathological recommencing of the loop and driving further damage. Colch treatment as much as the phospho-mimic S3E mutation blunts this feed-forward mechanism by correcting all pertinent elements of the pathway, ultimately leading to cardioprotection, even in a dystrophic environment produced by dystrophin loss. Contrary to S3E, the phospho-dead S3A mutation blunts the beneficial effect triggered by Colch. This is observed in dystrophic contexts and in a WT context, where dystrophin is expressed. Thus, the pathology caused by the S3A mutation is not restricted to DMD.

In a prior study (9), we challenged the pS-Cx43 mutations within mdx contexts. Although genetically identical to human DMD patients, the phenotype in mice is milder than in humans, in part because the dystrophin homologue utrophin compensates to a greater extent for the absence of dystrophin in mice. In this context, the S3E mutation led to protection against arrhythmias and long-term cardiac fibrosis, although an effect on survival under baseline conditions was not possible to assess. In an effort to study the full potency of our findings, we challenged pS-Cx43 mutations in more severe models of DMD. To this end, we incorporated the S3A and S3E mutations into mdx mice with partial and complete loss of utrophin (mdxutr+/- and mdxutr, respectively). Our current findings in these more severe dystrophic models also validate the protective effects of S3E against arrhythmias and cardiac fibrosis. Importantly, our current findings also demonstrate that the S3E mutation protects mice against early lethality. The effect is more appreciable in the mdxutr+/- than in mdxutr, where the differences are more robust. Thus, by S3E mutation, the Cx43/MT can bypass the absence of dystrophin and utrophin.

Since being established by Giepmans et. al (20, 35), the molecular partnership between Cx43 and MTs has gained significant interest in diverse pathologic states. Relevant to this study, various non-canonical roles of Cx43 modulating MTs have been demonstrated in cardiac pathophysiology (36). Internally translated Cx43 isoform GJA1-20k has been shown to regulate MT-based mitochondrial (37, 38), and Cx43 trafficking (22), actively participating towards cardioprotection. Furthermore, voltage-gated sodium channel Na_v_1.5 depends on Cx43 for proper forward MT-based trafficking to the plasma membrane (39). In the dystrophic heart, genetic inhibition of excessive activity of oxidized Ca^2+^/calmodulin Protein Kinase IIδ (Ox-CaMKII) has been shown to prevent ventricular arrhythmias (11). pS-Cx43 has been suggested to improve Cx43 assembly immediately before insertion into the ID, but its role in mediating cardiac MT trafficking remained undefined until this study (13, 40).

This study demonstrates that the interaction between Cx43 and β-tubulin can be improved in dystrophic hearts when pS-Cx43 is promoted, either directly through S3E mutation or indirectly through arresting MT growth. The Cx43–β-tubulin interaction was also improved in non-DMD human cardiac tissue compared to DMD, strengthening the clinical relevance of this mechanism. It has been suggested that enhancing the Cx43–β-tubulin interaction causes Cx43 to adhere to microtubules, promoting proper directionality as they reach the cell surface together, which may explain the improved directionality of mdxS3E MTs (41, 42). Alternatively, the excess of MTs in dystrophic hearts in physiological conditions may create trafficking routes that may either fail to tether properly at the ID or become re-routed to lateral sites, where the MT-Cx43 interaction may be more short-lived due to a lack of accompanying trafficking partners (42, 43). If excess MTs do not form, as in mdxS3E myocytes, a more stable Cx43–β-tubulin interaction may ensue, particularly right near the plasma membrane where pS-Cx43 occurs (13). The reduction of MTs may improve the stability and longevity of Cx43 at the gap junction since it has been suggested that MTs are involved in the degradation of Cx43 by trafficking endosomal vesicles (44).

This study has limitations. Although lateralized Cx43 hemichannels have been attributed as the primary mechanism for Ethidium uptake (14, 17, 42, 45), there is a possibility that S3A introduction also contributes to sarcolemma fragility of a dystrophic cardiomyocyte. Thus, we cannot formally rule out that a component of the dye is taken up not through the Cx43-S3A hemichannel directly but rather through other mechanisms that nevertheless depend on the presence of Cx43-S3A.

The study focuses on the heart of dystrophic mice, but the mutant Cx43 (S3E and S3A) is introduced globally. Thus, we cannot rule out changes in Cx43 remodeling, or Cx43 activity, in other tissues that may indirectly affect the outcome in the heart. However, we had reported that global mdxS3E and mdxS3A do not change the pathophysiology of the dystrophic skeletal muscle (9). Likewise, Colch treatment may have immunomodulatory activities, independent of its role as MT modulator (46).

It has been reported that the phospho-dead S3A promotes remodeling in WT contexts (40). Accordingly, in this study, we observe increased lateralization of WTS3A relative to WT. This highlights an additional mechanism of action that places phospho-dead Cx43 before the development of disarrayed MTs. Further studies need to be conducted to determine the contribution of dystrophin and non-dystrophic contexts to the disruptive capacity of S3A. Importantly, our current studies found that S3A blunts the beneficial effect of Colch in dystrophic cardiomyocytes.

MTs and Cx43 interaction is enhanced in the hearts of non-dystrophic versus dystrophic samples. This also applies to human non-DMD hearts, but a contribution to β-tubulin from non-myocytes, for example, fibroblasts, may interfere and obscure the outcome. However, we showed that the primary source of Cx43 in the heart is the myocyte - albeit in a mouse study (47). Since this study involves Cx43 immunoprecipitation of the complexes that are likely formed in-vivo and not after cell lysis, the contribution of β-tubulin from Cx43 non-expressing cells may not significantly count in the assay. Thus, the study likely reports an increase in MTs/Cx43 direct interaction in myocytes only.

In summary, these findings unveil a novel link between Cx43-phosphorylation and microtubule dynamics in dystrophic cardiomyocytes, with consequences in arrhythmias, cardiac fibrosis, and survival. Importantly, this regulation provides evidence that could be utilized in numerous ways to help develop therapeutic approaches for DMD cardiomyopathy. In the future, it may be appealing to test whether this regulation between MTs and Cx43 is operative only in dystrophic contexts, or the paradigm can be generalized to other systems where Cx43 is hypophosphorylated/remodeled.

## METHODS

### Mouse studies

Cx43 phospho-mutant (mdx:Cx43^S3A/S3A^, mdx:Cx43^S3E/S3E^) and Cx43-WT mdx (mdx:Cx43^+/+^) mice were generated as described in (9). WT and mdx control littermates and Cx43-mutant mice were analyzed at time points of 4-6 months of age. No differences were found between sexes in all the experiments performed in the study (6). Mice heterozygous for utrophin mdx:utr^**+/-**^ were bred to obtain mdx:utr^**+/-**^ and mdx:utr^null^ mice. Mdx:utr^**+/-**^ and mdx:Cx43^S3A/S3A^ or mdx:Cx43^S3E/S3E^ mice were crossed and genotyped as previously described in (9, 48), to obtain mdx:utr^**+/-**^:Cx43^S3E/S3E^ and mdx:utr^null^:Cx43^S3E/S3E^ littermates and their S3A counterparts. WT, mdx, mdxutr^**+/-**^, and Cx43-mutant mdxutr^**+/-**^ mice were analyzed at 10-14 months-old. Mdxutr^null^ and Cx43-mutant mdxutr^null^ mice were analyzed at 3-4 months-old. Sizes of experimental groups were based upon our prior studies (6, 9). See Supplemental Methods for details.

### Western Blotting

Snap frozen mouse, and human ventricular tissues were homogenized in either RIPA lysis buffer or Triton X-100 lysis buffer and processed as described in (9). See Supplemental Methods for details.

### Tissue Immunofluorescence and Cx43 Quantification

Immunofluorescent staining was performed as described previously (9). See Supplemental Methods for details

### Isolated Heart Ethidium Bromide Perfusion and Dye Uptake Quantification

Following Iso (5mg/kg) or vehicle injection, mice were treated with heparin (5000 U/kg) and anesthetized followed by cervical dislocation. Ethidium bromide (5μmol/L) was perfused into isolated hearts, then washed, fixed, sectioned, stained, and imaged as described in (17). See Supplemental Methods for details on quantifications.

### Ventricular Cardiomyocyte Isolation

Single ventricular cardiomyocytes were isolated from mouse hearts using a Langendorff perfusion system as described in (9) and then were incubated in the presence or absence of Colchicine (10 μM) for 1 hour. Cells were kept in isolation buffer before fixation for immunofluorescent analyses.

### Immunofluorescence of Isolated Cardiomyocytes

Freshly isolated cardiomyocytes were plated on laminin-coated (10μg/mL) chamber slides, fixed in 4% paraformaldehyde, washed in PBS, and permeabilized with 0.5% Triton X-100 in PBS. Cells were incubated in blocking buffer, then with β-tubulin (Sigma T8328, 1:1000, mouse) in blocking buffer overnight and finally with Alexa Fluor secondary antibodies. Confocal Z-stacks of a step size of 0.5μm thickness were acquired at 60x magnification on an Olympus Fluoview 1000 Confocal Laser Scanning Microscope, then processed as the sum of 15 images from the inter-myofibrillar region (> 3 μm from surface (19)) of the cardiomyocyte. Gray-scale, 32-bit images were subject to contrast enhancement using Adobe Photoshop for figure presentation only. See Supplemental Methods for details.

### Confocal Microtubule Directionality and Density

Confocal images of cardiomyocyte β-tubulin processed above were converted to 8-bit images, oriented horizontally, and analyzed using the texture direction technique (TeDT, software available upon request to Dr. Ralston) (23). Selections were made as large as possible, excluding areas around nuclei and the edge of the cell. Microtubule density was measured by converting the ROI from TeDT analysis into a binary image and quantified using ImageJ software, with % area fraction as a final readout. A two-way ANOVA was used to assess the statistical difference between the directionality histograms of all mouse models used for analysis. A one-way ANOVA with Tukey’s post-hoc test was used to assess the statistical differences between the groups in microtubule density.

### Mini Osmotic Pump Implantation

Mice randomly chosen for osmotic pump implantation were weighed and anesthetized with isoflurane. Osmotic pumps (Alzet model 1004, 28-day) were inserted through a small incision between the scapulae, and the incision was then closed with wound clips. Osmotic pumps contained either Saline (vehicle) or Colchicine (Sigma C9754, dissolved in Saline, 0.4mg/kg/day) and remained in mice for 28 days. At the end of the study, mice were euthanized, and cardiac tissue was collected for downstream immunohistochemical and biochemical assays.

### Telemetry Device Implantation

A Data Sciences International (DSI) telemetry transmitter to measure ECG was implanted as follows: After the animal is anesthetized by isoflurane, shaved, and aseptically prepared, a 1-2 cm transverse incision was made in the left inguinal area. A subcutaneous pocket was created on the left side of the abdomen toward the caudal edge of the rib cage. The telemetric transmitter, a tubular disk 1 cm in diameter X 2 cm length, was inserted into the subcutaneous pocket and secured to the abdominal wall with sutures in three places on the left flank. Two wires for the ECG lead are tunneled subcutaneously; the ends are stripped to provide transmission and then coiled and secured to the underlying muscle tissue. The positive ECG lead wire was tunneled to the lower left rib cage and fixed to the abdominal muscle; the negative ECG lead is tunneled to the caudal end of the right scapula and fixed to the pectoralis muscle on the right side. Finally, the transverse inguinal incision is closed in two layers using interrupted mattress sutures and 3-0 nylon. This procedure does not require externalization of the wires, thus minimizing the possibility of infection and allowing the animal to be monitored from the outside without interference.

### Isoproterenol Stress Test Recording

Once recovered from telemetry device implantation surgery (3-5 days), mice were subjected to a 3-day Isoproterenol stress study. Mice were first weighed and separated into single cages that were placed on telemetry receivers. A 1-hour baseline reading was taken to monitor activity and ECG readings. Next, mice were intraperitoneally injected with Isoproterenol (Iso, Sigma I6504, 5mg/kg) and then again 24 and 48 hours afterward. Mice were constantly recorded and observed for changes in activity and morbidity. Electrocardiographic data was analyzed by Lab Chart 8 software (Life Science Data Acquisition Software).

### Immunoprecipitation

1 mg of Triton-soluble samples prepared as described in (9) were incubated with 4 μl of Cx43 antibody (Sigma C6219, rabbit) and subsequently immunoprecipitated. See Supplemental Methods for details.

### Fibrosis Staining and Quantification

Masson Trichrome staining and quantification were performed as described previously (7).

### Human Samples

Three non-DMD and three DMD male human heart samples were obtained from the University of Maryland Brain and Tissue Bank, a member of the NIH NeuroBioBank network. All samples were dissected post-mortem. DMD1 cause of death was attributable to cardiac failure at age 15, DMD2 cause of death was pulmonary thromboembolism at age 17, and DMD3 cause of death unknown. Informed consent was obtained from all subjects from whom tissues were analyzed.

### Statistics

All results are presented as mean ± standard error of the mean (SEM). Statistical analyses were performed using parametric analysis in GraphPad Prism software. Statistical significance between multiple groups was analyzed by one-way ANOVA parametric testing followed by Tukey’s multiple comparisons test. Statistical significance amongst mice with osmotic pumps and were ISO injected was analyzed by two-way ANOVA followed by Tukey’s multiple comparisons test. In the case of two groups, we performed paired, two-tailed t-tests. P values less than 0.05 were considered significant for all statistical tests. Representative p values and symbols are described in figure legends. Most experiments and analyses of endpoint readouts were carried out in a blinded fashion.

### Study approval

All animal experiments were approved by the IACUCs of Rutgers New Jersey Medical School and Baylor College of Medicine and performed in accordance with NIH guidelines. non-DMD and DMD male human heart samples were obtained from the University of Maryland Brain and Tissue Bank, a member of the NIH NeuroBioBank network. All samples were dissected post-mortem. All human experiments were approved by the IRB of Rutgers University and performed following relevant guidelines and regulations. Informed consent was obtained for all subjects from whom tissues were analyzed.

## ACKNOWLEDGEMENTS

We thank Dr. Glenn I. Fishman (New York University) for providing the original S3A and S3E founder mice for our colonies; Dr. Paul D. Lampe (Fred Hutchison Cancer Center) for providing custom Cx43-pS325/S328/S330 antibody; Drs. Wenhua Liu and Evelyn Ralston (Light Image Section, National Institute of Arthritis and Musculoskeletal and Skin Diseases, NIH) for providing the TeDT analysis program; Delong Zhou and Halle Sarkodie for lab assistance; and Luke Fritzy (Rutgers NJMS) for help with histology.

## AUTHOR CONTRIBUTIONS

Conceived and designed the experiments: EH, MAL, DF, JEC, JN, NS. Performed the experiments: EH, JN, MAL, AC. Analyzed the data: EH, JN, MAL, DF, JEC, LHX, NS, GGR XHTW. Contributed reagents/materials/analysis tools: DF, JEC, NS, GGR. Wrote the paper: EH, JN. All authors reviewed and approved the final draft.

## REFERENCES

1. Birnkrant DJ, et al. Diagnosis and management of Duchenne muscular dystrophy, part 2: respiratory, cardiac, bone health, and orthopaedic management. Lancet Neurol. 2018;17(4):347–61.

2. Kamdar F, and Garry DJ. Dystrophin-Deficient Cardiomyopathy. J Am Coll Cardiol. 2016;67(21):2533–46.

3. Lapidos KA, et al. The dystrophin glycoprotein complex: signaling strength and integrity for the sarcolemma. Circ Res. 2004;94(8):1023–31.

4. Meyers TA, and Townsend D. Cardiac Pathophysiology and the Future of Cardiac Therapies in Duchenne Muscular Dystrophy. International Journal of Molecular Sciences. 2019;20(17):4098.

5. Shirokova N, and Niggli E. Cardiac phenotype of Duchenne Muscular Dystrophy: Insights from cellular studies. Journal of Molecular and Cellular Cardiology. 2013;58:217–24.

6. Gonzalez JP, et al. Selective Connexin43 Inhibition Prevents Isoproterenol-Induced Arrhythmias and Lethality in Muscular Dystrophy Mice. Scientific Reports. 2015;5:13490.

7. Gonzalez JP, et al. Small Fractions of Muscular Dystrophy Embryonic Stem Cells Yield Severe Cardiac and Skeletal Muscle Defects in Adult Mouse Chimeras. Stem Cells (Dayton, Ohio). 2016;35(3):597–610.

8. Gonzalez JP, et al. Normalization of connexin 43 protein levels prevents cellular and functional signs of dystrophic cardiomyopathy in mice. Neuromuscular Disorders. 2018;28(4):361–72.

9. Himelman E, et al. Prevention of connexin-43 remodeling protects against Duchenne muscular dystrophy cardiomyopathy. The Journal of Clinical Investigation. 2020;130(4):1713–27.

10. Wang Q, et al. Oxidized CaMKII (Ca2+/Calmodulin-Dependent Protein Kinase II) Is Essential for Ventricular Arrhythmia in a Mouse Model of Duchenne Muscular Dystrophy. Circulation: Arrhythmia and Electrophysiology. 2018;11(4):e005682.

11. Erickson JR, et al. A Dynamic Pathway for Calcium-Independent Activation of CaMKII by Methionine Oxidation. Cell. 2008;133(3):462–74.

12. Cooper CD, and Lampe PD. Casein kinase 1 regulates connexin-43 gap junction assembly. J Biol Chem. 2002;277(47):44962–8.

13. Lampe PD, et al. Analysis of Connexin43 phosphorylated at S325, S328 and S330 in normoxic and ischemic heart. Journal of Cell Science. 2006;119(16):3435–42.

14. Contreras JE, et al. Gating and regulation of connexin 43 (Cx43) hemichannels. Proc Natl Acad Sci U S A. 2003;100(20):11388–93.

15. Johnson RG, et al. Connexin Hemichannels: Methods for Dye Uptake and Leakage. J Membr Biol. 2016;249(6):713–41.

16. Figueroa XF, et al. Diffusion of nitric oxide across cell membranes of the vascular wall requires specific connexin-based channels. Neuropharmacology. 2013;75:471–8.

17. Lillo MA, et al. S-nitrosylation of connexin43 hemichannels elicits cardiac stress–induced arrhythmias in Duchenne muscular dystrophy mice. JCI Insight. 2019;4(24).

18. Zhao Z, et al. Overexpression of adenylyl cyclase type 5 (AC5) confers a proarrhythmic substrate to the heart. American Journal of Physiology-Heart and Circulatory Physiology. 2014;308(3):H240–H9.

19. Loehr JA, et al. NADPH oxidase mediates microtubule alterations and diaphragm dysfunction in dystrophic mice. Elife. 2018;7:e31732.

20. Giepmans BNG, et al. Gap junction protein connexin-43 interacts directly with microtubules. Current Biology. 2001;11(17):1364–8.

21. Chkourko HS, et al. Experimental: Remodeling of mechanical junctions and of microtubule-associated proteins accompany cardiac connexin43 lateralization. Heart Rhythm. 2012;9:1133–40.e6.

22. Basheer WA, et al. GJA1-20k Arranges Actin to Guide Cx43 Delivery to Cardiac Intercalated Discs. Circ Res. 2017;121(9):1069–80.

23. Liu W, and Ralston E. A new directionality tool for assessing microtubule pattern alterations. Cytoskeleton (Hoboken, NJ). 2014;71(4):230–40.

24. Caporizzo MA, et al. Cardiac microtubules in health and heart disease. Experimental Biology and Medicine. 2019;244(15):1255–72.

25. Strakova J, et al. Dystrobrevin increases dystrophin’s binding to the dystrophin–glycoprotein complex and provides protection during cardiac stress. Journal of Molecular and Cellular Cardiology. 2014;76:106–15.

26. Prosser BL, et al. X-ROS Signaling: Rapid Mechano-Chemo Transduction in Heart. Science. 2011;333(6048):1440–5.

27. Khairallah RJ, et al. Microtubules Underlie Dysfunction in Duchenne Muscular Dystrophy. Science Signaling. 2012;5(236):ra56–ra.

28. Giepmans BN. Gap junctions and connexin-interacting proteins. Cardiovasc Res. 2004;62(2):233–45.

29. Janssen PM, et al. Utrophin deficiency worsens cardiac contractile dysfunction present in dystrophin-deficient mdx mice. Am J Physiol Heart Circ Physiol. 2005;289(6):H2373–8.

30. Kelly MG, et al. Skeletal muscle fibrosis in the mdx/utrn+/-mouse validates its suitability as a murine model of Duchenne muscular dystrophy. PLoS ONE, Vol 10, Iss 1, p e0117306 (2015). (1):e0117306.

31. Yucel N, et al. Humanizing the mdx mouse model of DMD: the long and the short of it. NPJ Regenerative medicine. 2018;3(1):1–11.

32. Gutpell KM, et al. Skeletal muscle fibrosis in the mdx/utrn+/-mouse validates its suitability as a murine model of Duchenne muscular dystrophy. PLoS One. 2015;10(1):e0117306.

33. Delfín DA, et al. Cardiomyopathy in the dystrophin/utrophin-deficient mouse model of severe muscular dystrophy is characterized by dysregulation of matrix metalloproteinases. Neuromuscular Disorders. 2012;22:1006–14.

34. Spurney CF. Cardiomyopathy of duchenne muscular dystrophy: Current understanding and future directions. Muscle & Nerve. 2011;44(1):8–19.

35. Giepmans BN, et al. Connexin-43 interactions with ZO-1 and alpha- and beta-tubulin. Cell Commun Adhes. 2001;8(4-6):219–23.

36. Rusiecka OM, et al. Canonical and Non-Canonical Roles of Connexin43 in Cardioprotection. Biomolecules. 2020;10(9):1225.

37. Basheer WA, et al. Stress response protein GJA1-20k promotes mitochondrial biogenesis, metabolic quiescence, and cardioprotection against ischemia/reperfusion injury. JCI Insight. 2018;3(20).

38. Fu Y, et al. Cx43 Isoform GJA1-20k Promotes Microtubule Dependent Mitochondrial Transport. Frontiers in physiology. 2017;8:905–.

39. Agullo-Pascual E, et al. Super-resolution imaging reveals that loss of the C-terminus of connexin43 limits microtubule plus-end capture and NaV1.5 localization at the intercalated disc. Cardiovascular Research. 2014;104(2):371–81.

40. Remo BF, et al. Phosphatase-resistant gap junctions inhibit pathological remodeling and prevent arrhythmias. Circ Res. 2011;108(12):1459–66.

41. Francis R, et al. Connexin43 Modulates Cell Polarity and Directional Cell Migration by Regulating Microtubule Dynamics. PLOS ONE. 2011;6(10):e26379.

42. Rhett JM, et al. Mechanism of action of the anti-inflammatory connexin43 mimetic peptide JM2. Am J Physiol Cell Physiol. 2017;313(3):C314–c26.

43. Smyth JW, et al. Limited forward trafficking of connexin 43 reduces cell-cell coupling in stressed human and mouse myocardium. The Journal of Clinical Investigation. 2010;120(1):266–79.

44. Guan X, and Ruch RJ. Gap junction endocytosis and lysosomal degradation of connexin43-P2 in WB-F344 rat liver epithelial cells treated with DDT and lindane. Carcinogenesis. 1996;17(9):1791–8.

45. De Maio A, et al. Gap junctions, homeostasis, and injury. Journal of Cellular Physiology. 2002;191(3):269–82.

46. Leung YY, et al. Colchicine—Update on mechanisms of action and therapeutic uses. Seminars in Arthritis and Rheumatism. 2015;45:341–50.

47. Nouet J, et al. Connexin-43 reduction prevents muscle defects in a mouse model of manifesting Duchenne muscular dystrophy female carriers. Sci Rep. 2020;10(1):5683.

48. Gonzalez JP, et al. Neuronal nitric oxide synthase localizes to utrophin expressing intercalated discs and stabilizes their structural integrity. Neuromuscul Disord. 2015;25(12):964–76.

