## Supplemental material for "Microtubule-Connexin-43 regulation suppresses arrhythmias and fibrosis in Duchenne muscular dystrophy mice"

### Supplemental Methods

*Mouse studies*

WT (C57BL/6J) mice were purchased from Jackson Laboratories (Bar Harbor, ME). Cx43 phospho-mutant (mdx:Cx43$^{S3A/S3A}$, mdx:Cx43$^{S3E/S3E}$) and Cx43-WT mdx (mdx:Cx43$^{+/+}$) mice were generated, back-crossed 7-10 generations and genotyped as described in (1). WT and mdx control littermates and Cx43-mutant mice were analyzed at time points of 4-6 months of age. No differences were found between the sexes in all the experiments performed in the study (2). Mice heterozygous for utrophin mdx:utr$^{+/-}$ were bred to obtain mdx:utr$^{+/-}$ and mdx:utr$^{null}$ mice. Mdx:utr$^{+/-}$ and mdx:Cx43$^{S3A/S3A}$ or mdx:Cx43$^{S3E/S3E}$ mice were crossed. Heterozygous progeny, mdx:utr$^{+/-}$:Cx43$^{S3E/WT}$ or mdx:utr$^{+/-}$:Cx43$^{S3A/WT}$, were crossed to yield mdx:utr$^{+/-}$:Cx43$^{S3E/S3E}$ or mdx:utr$^{+/-}$:Cx43$^{S3A/S3A}$. These were backcrossed at least six generations and genotyped as previously described in (1, 3), to obtain mdx:utr$^{+/-}$:Cx43$^{S3E/S3E}$ and mdx:utr$^{null}$:Cx43$^{S3E/S3E}$ littermates and their S3A counterparts. WT, mdx, mdxutr$^{+/-}$, and Cx43-mutant mdxutr$^{+/-}$ mice were analyzed at 10-14 months-old. Mdxutr$^{null}$ and Cx43-mutant mdxutr$^{null}$ mice were analyzed at 3-4 months-old. All mice were maintained on a 12-hour light-dark cycle with free access to food and water. Sizes of experimental groups were based upon our prior studies (1, 2).

*Western Blotting*

Snap frozen mouse and human ventricular tissues were homogenized in either RIPA lysis buffer or Triton X-100 lysis buffer and processed as described in (1). Membranes were incubated with either Cx43 (Sigma C6219, 1:10000, rabbit), β-tubulin (Sigma T8328, 1:1000, mouse), pS325/S328/S330-Cx43 (Provided by Dr. Paul Lampe, 1:1000, mouse), , or Vinculin (Sigma V9131, 1:2000, mouse, loading control). Protein band densities were visualized with ImageLab software (Bio-Rad) and quantified by Fiji ImageJ.

*Tissue Immunofluorescence and Cx43 Quantification*

Immunofluorescent staining was performed on cryo-embedded cardiac cryosections with Cx43 (Sigma C6219, 1:2000, rabbit) and N-Cadherin (Invitrogen 33-3900, 1:400, mouse) as described in (1). Confocal Z-stacks of a step size of 0.5μm thickness (approximately 12 slices per image) were acquired at 60x magnification on an Olympus Fluoview 1000 Confocal Laser Scanning Microscope using the Fluoview software. The Cx43/N-Cadherin images were separated into separate channels and processed as maximum intensity z-stack projections in Fiji before analysis. Maximum intensity z-stacks were processed, and Cx43 intensities at N-Cadherin positive intercalated discs were calculated as described in (1).

*Isolated Heart Ethidium Bromide Perfusion and Dye Uptake Quantification*

Following Iso (5mg/kg) or vehicle injection, mice were treated with heparin (5000 U/kg) and anesthetized with Avertin (290mg/kg). Ethidium bromide (5μmol/L) was perfused into isolated hearts, washed, fixed, sectioned, stained, and imaged as described in (4). Slides were imaged on a Nikon Eclipse T microscope. To quantify relative Ethidium uptake, images taken at the same exposure settings were processed in ImageJ for ethidium red autofluorescence (555nm excitation/580nm emission) and DAPI (345nm excitation/455nm emission) nuclei. In a blinded fashion, the ratio of DAPI to Ethidium signal for each nucleus was quantified and expressed as arbitrary units (AU) using ImageJ (NIH), as performed in (1, 4).

*Immunofluorescence of Isolated Cardiomyocytes*

Freshly isolated cardiomyocytes were plated on laminin-coated (10μg/mL) chamber slides and allowed to adhere for 30 minutes at room temperature. Cells were fixed in 4% paraformaldehyde for 15 minutes at room temperature, washed 3 x 5 minutes in PBS, and permeabilized with 0.5% Triton X-100 in PBS for 20 minutes. Following 3 x 5 minute washes in PBS, cells were incubated in blocking buffer (2% NGS, 2% BSA, 0.3M glycine in PBS) for 1 hour at room temperature. Then, cells were incubated with β-tubulin (Sigma T8328, 1:1000, mouse) in blocking buffer overnight at 4ºC. Following three washes in PBS, cells were incubated for an hour at room temperature with Alexa Fluor secondary antibodies (Invitrogen) in blocking buffer (1:250). Slides were subsequently washed in PBS, and coverslips were mounted using ProLong Gold antifade reagent containing DAPI. Confocal Z-stacks of a step size of 0.5μm thickness were acquired at 60x magnification on an Olympus Fluoview 1000 Confocal Laser Scanning Microscope using the Fluoview software. The β-tubulin images were then processed as the sum of 15 images from the inter-myofibrillar region ( > 3 μm from surface (5)) of the cardiomyocyte. Gray-scale, 32-bit images were subject to contrast enhancement using Adobe Photoshop for figure presentation only.

*Immunoprecipitation*

1 mg of Triton-soluble samples prepared as described in (1) were incubated with 4 μl of Cx43 antibody (Sigma C6219, rabbit) and rotated end to end for 6 hours at 4°C. Next, antigen-antibody mixtures were added to 50μl of pre-washed Protein A/G Magnetic Beads (Invitrogen) and incubated for 16 hours on end-to-end rotation at 4°C. The beads were washed three times in 1mL of cold 1% Triton X-100 lysis buffer supplemented with protease and phosphatase inhibitors, then washed in 1mL of cold PBS. The beads were then magnetized, and the supernatant discarded. 80μl of 2x SDS sample buffer was added to the beads, gently mixed, then heated at 96°C for 10 minutes. The supernatants containing the final IP product were extracted from the beads by magnetization and run on 10% SDS-PAGE gels as described in the "Western Blotting" section. Either rabbit anti-mouse or mouse anti-rabbit IgG Light Chain Specific HRP monoclonal antibodies (Cell Signaling 58802S or 45262S, 1:2000) were added to the membranes in blocking solution for 2 hours at room temperature to avoid detection of denatured mouse or rabbit IgG heavy chains of the primary antibody in immunoprecipitated samples. The membranes were subsequently washed and imaged as described in the "Western Blotting" section.

*Fibrosis Staining and Quantification*

Masson Trichrome staining and quantification were performed as described previously(6) and processed using ImageJ software. Fibrosis calculations used the blue-stain area divided by the tissue's total area for 3-4 images at 4x and 10x magnification.

**Supplemental Figures**

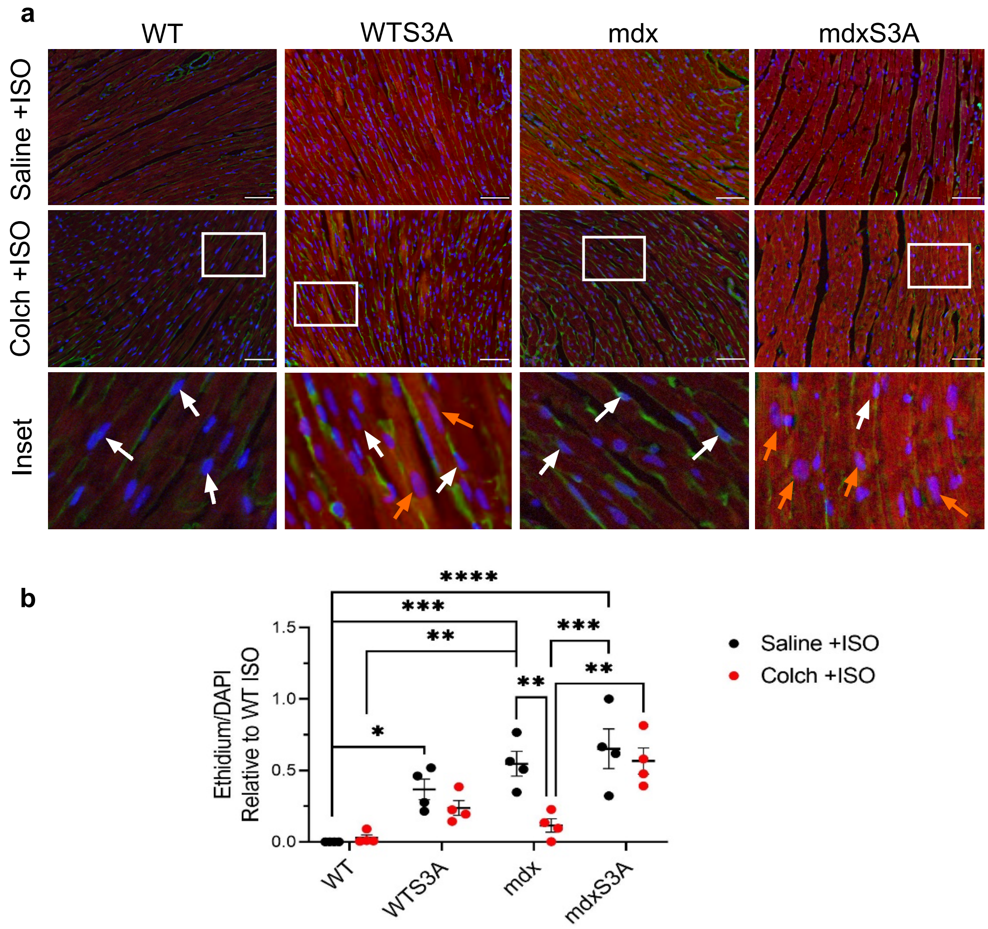


**Supplemental Figure S1**. Aberrant Ethidium uptake in the hearts of mdxS3A mice following Colch treatment. (**a**) Representative immunofluorescence images of Ethidium uptake of heart cryosections after perfusion with Ethidium (5μM) following 4-week Saline (Saline +ISO) or Colchicine (Colch +ISO) treatment and an acute injection of ISO (5mg/kg), processed in Fiji ImageJ. Cryosections were visualized for Ethidium (red) and stained for WGA (green) and nuclei (DAPI, blue). White boxes indicate areas magnified in the bottom row (insets). White arrows indicate nuclei that do not express Ethidium; orange arrows indicate nuclei positive for Ethidium stain. Scale bar, 50μm. (**b**) Quantification of the dye uptake in ISO-stressed hearts following Saline (Saline +ISO) or Colch (Colch +ISO) treatment. *n* = 4 per group per treatment. (2-way ANOVA, Tukey's multiple comparisons test, **** = $p < 0.0001$, *** = $p < 0.001$, ** = $p < 0.01$, * = $p < 0.05$) Data are presented as means ± SEM.

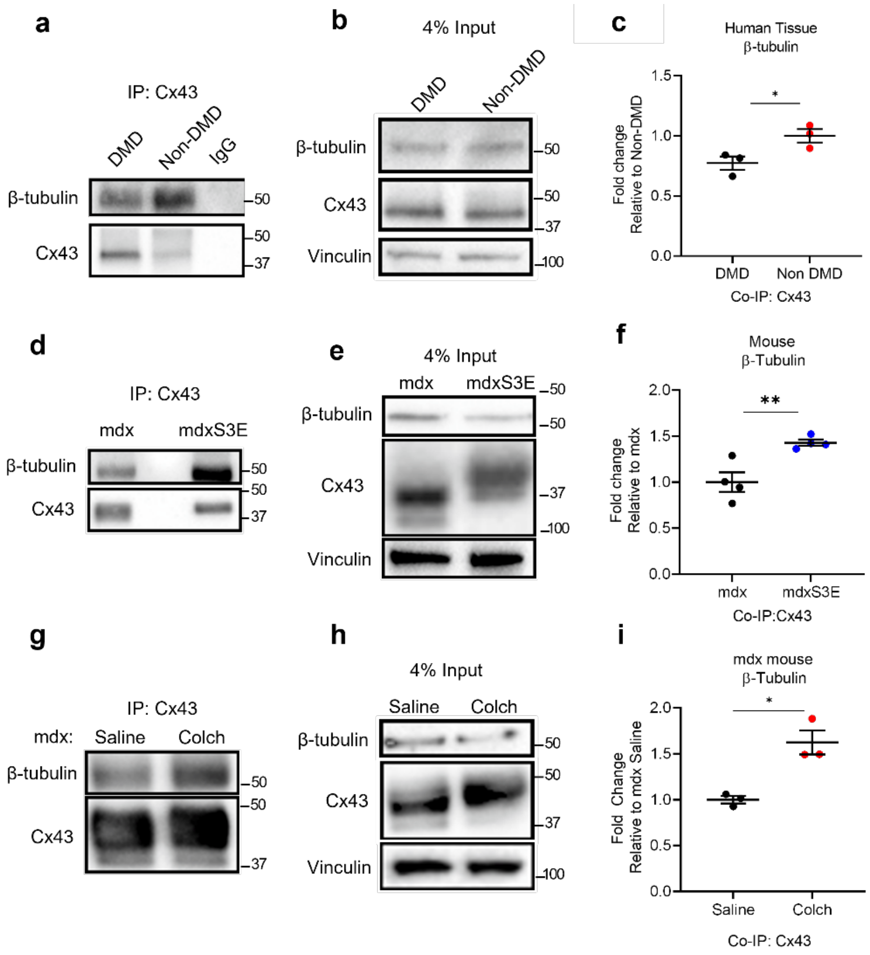


**Supplemental Figure S2**. Improved Cx43-β-tubulin interaction in non-DMD human cardiac lysates. (**a**) Representative results of a co-immunoprecipitation (co-IP) assay in DMD and non-DMD human cardiac lysates. (**b**) Representative results of a western blot performed with 4% of DMD and non-DMD human lysates used for immunoprecipitation, probed for β-tubulin, Cx43, and Vinculin (loading control). (**c**) Quantification of β-tubulin expressed following co-IP in (**a**), normalized to input loading levels, expressed as fold change relative to non-DMD. *n*=3 per group. (Two-sided t-test, t=1.641, df=4,*=p<0.05). (**d**) Representative results of a co-IP assay using Cx43 antibody, followed by β-tubulin and Cx43 western blotting in mdx and mdxS3E lysates. (**e**) Representative results of western blot performed with 4% of mdx and mdxS3E lysates used for immunoprecipitation, probed for β-tubulin, Cx43, and Vinculin (loading control). (**f**) Quantification of β-tubulin expressed following co-IP in (**d**), normalized to input loading levels, expressed as fold change relative to mdx. *n*=4 both groups. (Two-sided t-test, t=3.785, df=6, **=p<0.01). (**g**) Representative results of a co-IP assay described in (**d**) in Saline and Colch-treated mdx lysates. (**h**) Representative results of western blot performed with 4% of Saline and Colch-treated mdx lysates used for immunoprecipitation, probed for β-tubulin, Cx43, and Vinculin (loading control). (**i**) Quantification of β-tubulin expressed following co-IP in (**g**), normalized to input loading levels, expressed as fold change relative to mdx saline. *n*=3 both groups. (Two-sided t-test, t=4.597, df=4, *=p<0.05).

**Supplemental References**


1. Himelman E, Lillo MA, Nouet J, Gonzalez JP, Zhao Q, Xie L-H, et al. Prevention of connexin-43 remodeling protects against Duchenne muscular dystrophy cardiomyopathy. *The Journal of Clinical Investigation.* 2020;130(4):1713-27.
2. Gonzalez JP, Ramachandran J, Xie L-H, Contreras JE, and Fraidenraich D. Selective Connexin43 Inhibition Prevents Isoproterenol-Induced Arrhythmias and Lethality in Muscular Dystrophy Mice. *Scientific Reports.* 2015;5:13490.
3. Gonzalez JP, Crassous PA, Schneider JS, Beuve A, and Fraidenraich D. Neuronal nitric oxide synthase localizes to utrophin expressing intercalated discs and stabilizes their structural integrity. *Neuromuscul Disord.* 2015;25(12):964-76.
4. Lillo MA, Himelman E, Shirokova N, Xie L-H, Fraidenraich D, and Contreras JE. S-nitrosylation of connexin43 hemichannels elicits cardiac stress–induced arrhythmias in Duchenne muscular dystrophy mice. *JCI Insight.* 2019;4(24).
5. Loehr JA, Wang S, Cully TR, Pal R, Larina IV, Larin KV, et al. NADPH oxidase mediates microtubule alterations and diaphragm dysfunction in dystrophic mice. *Elife.* 2018;7:e31732.
6. Gonzalez JP, Kyrychenko S, Kyrychenko V, Schneider JS, Granier CJ, Himelman E, et al. Small Fractions of Muscular Dystrophy Embryonic Stem Cells Yield Severe Cardiac and Skeletal Muscle Defects in Adult Mouse Chimeras. *Stem Cells (Dayton, Ohio).* 2016;35(3):597-610.
